# Selection for increased post-infection survival ameliorates mating induced immune suppression in *Drosophila melanogaster* females

**DOI:** 10.1101/2022.07.01.498387

**Authors:** Aabeer Basu, Aparajita Singh, B G Ruchitha, Nagaraj Guru Prasad

## Abstract

Sexual activity (mating) negatively affects immune function in various insect species, in both sexes. In the experiments reported in this manuscript, we tested if hosts adapted to regular pathogen challenges are less susceptible to mating induced immune suppression, using experimentally evolved *Drosophila melanogaster* populations selected for increased post-infection survival when infected with a Gram-positive bacterium, *Enterococcus faecalis*. Mating increased susceptibility of females to bacterial pathogens, but in a pathogen specific manner. Mating-induced increase in susceptibility was also affected by host evolutionary history, with females from selected populations exhibiting similar post-infection survival irrespective of mating status, while females from control populations became more susceptible to bacterial infections after mating. Post-infection survival of males, irrespective of their evolutionary history, was not affected by their mating status. We therefore conclude that hosts evolved to better survive bacterial infections are also better at resisting mating-induced increase in susceptibility to infections in *Drosophila melanogaster*.

## 1. Introduction

Reproduction-immunity trade-offs in insects and other invertebrates work in either direction: infected hosts (while mounting an immune defense) exhibit reduced reproductive output, and hosts investing towards reproduction often have compromised immune function (Lawniczak et al 2007, Schwenke et al 2016). Sexual activity (mating) induced immune suppression has been observed in very many insect species, including *Matrona basilaris japonica* (Japanese calopterygid damselfly, Siva-Jothy et al 1998), *Tenebrio molitor* (mealworm beetle, Rolf and Siva-Jothy 2002), *Allonemobius socius* (striped ground cricket, Fedorka et al 2004), *Formica paralugubris* (wood ants, Castella et al 2009), and *Drosophila melanogaster* (McKean and Nunney 2001, McKean and Nunney 2005, Fedorka et al 2007, Short and Lazzaro 2010, Khan and Prasad 2011, Short et al 2012, Schwenke and Lazzaro 2017, Gupta et al 2021, Gordon et al 2022), in case of both males and females. Mating induced immune suppression can manifest as increased post-infection mortality, reduced capacity of clearing systemic pathogen load, and/or down regulation of a specific component of the immune system.

There are many nuances associated with mating induced immune suppression in insects. One, different components of the immune system may be affected differently, in terms of both direction and degree, by sexual activity in a particular insect species. For example, mating reduces hemocyte count, encapsulation ability, and lytic activity, but increases phenol oxidase (PO) activity in female crickets (*A. socius*, Fedorka et al 2004). Similarly in wood ant queens (*F. paralugubris*), mating reduces PO activity but increases antibacterial defenses (Castella et al 2009). Two, the same immune component may be affected differently in different insect species. For example, PO activity increases after mating in female crickets (*A. socius*, Fedorka et al 2004), but decreases after mating in case of female meal worm beetles (*T. molitor*, Rolf and Siva-Jothy 2002) and wood ant queens (*F. paralugubris*, Castella et al 2009). Three, post-mating immune suppression can be sex specific. For example, mating increases PO activity in females but reduces PO activity in male crickets (*A. socius*, Fedorka et al 2004).

Four, differences in individual components of immune system may not translate into actualized resistance to diseases, in terms of post-infection survival and systemic pathogen clearance. For example, in *D. melanogaster* ovoD1 mutant females (in which oogenesis is inhibited before vitellogenesis), mating leads to immediate upregulation of various anti-microbial peptide (AMP) genes, but mated females die more compared to virgin females when infected with the bacterium *Pseudomonas aeruginosa* (Fedorka et al 2007). Five, in cases where post-infection survival is affected by mating, the effects can be pathogen specific. For example, mated *D. melanogaster* females are more susceptible to infections with *Providencia rettgeri* and *Providencia alcalifaciens* compared to virgins, but not in case of infections with *Enterococcus faecalis* and *Pseudomonas entomophila* (Short and Lazzaro 2010). In case of all these four pathogens, the differences in survival between virgin and mated females was correlated with differences in systemic bacterial load (Short and Lazzaro 2010).

Six, in certain insect species, mating can have a positive effect on post-infection fitness of the host, in both females (reviewed in Oku et al 2019) and males. For example, mating reduces likelihood of infection by Trypanosoma parasite *Crithidia bombi* in bumblebee (*Bombus terrestris*) males and queens (Barribeau and Schmid-Hempel 2017). Mating also improves survival following bacterial infection in *D. melanogaster* males, but in a pathogen specific manner (Gupta et al 2013, Syed et al 2020). Various studies using *D. melanogaster* females have reported mating induced changes (mostly upregulation) in expression pattern of genes involved in immune defense, without measuring post-infection survival of the host or systemic pathogen clearance (McGraw et al 2004, Peng et al 2005, Winterhalter and Fedorka 2009, Gioti et al 2012, Fricke et al 2020). Since changes in sub-organismal immune components do not always translate into differential host survival (Adamo 2004b), it becomes difficult to interpret the results from such studies from the vantage point of eco-immunology (Adamo 2004a). And seven, observed effect of mating on the immune system is often dependent on the time elapsed since the mating event, for example as observed in *F. paralugubris* (Castella et al 2009) and *D. melanogaster* (Fedorka et al 2007, Winterhalter and Fedorka 2009, Short et al 2012; but see Gordon et al 2022).

In the present study we tested if host evolutionary history determined the difference between post-infection survival of virgin and mated flies in *Drosophila melanogaster*, in case of both females and males. We evolved a set of replicated fly populations, selecting for increased post-infection survival following infection with a Gram-positive bacterium, *Enterococcus faecalis*. The selected populations evolved better post infection survival compared to the ancestrally paired control populations within 35 generations of forward selection (Singh et al 2021). We subjected virgin and mated females from the selected and the control populations to infection with three pathogens: *E. faecalis*, the native pathogen used for selection, and two novel pathogens, *Bacillus thuringiensis* and *Pseudomonas entomophila*. Our results indicate that whether mated females die more following infection, compared to the virgin flies, is contingent upon host evolutionary history, pathogen identity, and the interaction between these two factors. Males, both mated and virgins, were infected only with the native pathogen. Results show that for this pathogen, mating does not change the susceptibility of males to infection.

## 2. Materials and Methods

### 2.1 Pathogen handling and infection protocol

Three bacterial pathogens were used in this study: (a) *Enterococcus faecalis* (Lazzaro et al 2006), a Gram-positive bacterium, which was used in regular maintenance of the EPN populations (Singh et al 2021), and in experiments, and (b) *Bacillus thuringiensis* (DSM 2046, obtained from Leibniz Institute DSMZ-German Collection of Microorganisms and Cell Cultures GmbH), a Gram-positive bacterium, and (c) *Pseudomonas entomophila* (strain L48, Vodovar et al 2005, Mulet et al 2012), a Gram-negative bacterium, both of which were used only for experimental infections.

The bacteria are stored as glycerol stocks (17%) in −80 °C. To obtain live bacterial cells for infections, 10 ml lysogeny broth (Luria Bertani Broth, Miler, HiMedia) is inoculated with glycerol stocks of the required bacterium and incubated overnight with aeration (150 rpm shaker incubator) at suitable temperature (37 °C for *E. faecalis*, 30 °C for *B. thuringiensis*, and 27 °C for *P. entomophila*). 100 microliters from this primary culture is inoculated into 10 ml fresh lysogeny broth and incubated for the necessary amount of time to obtain confluent (OD_600_= 1.0-1.2) cultures. The bacterial cells are pelleted down using centrifugation and resuspended in sterile MgSO_4_ (10 mM) buffer to obtain the required optical density (OD_600_) for infection. Flies are infected, under light CO_2_ anaesthesia, by pricking them on the dorsolateral side of their thorax with a 0.1 mm Minutien pin (Fine Scientific Tools, USA) dipped in the bacterial suspension. Sham-infections (injury controls) are carried out in the same fashion, except by dipping the pins in sterile MgSO_4_ (10 mM) buffer.

During regular maintenance of the EPN populations, the flies from the E_1-4_ populations (see below) are infected with an *E. faecalis* suspension of OD_600_ = 1.2. For all experimental infections (see below), for all three pathogens, flies were infected with a bacterial suspension of OD_600_ = 1.0.

### 2.2 EPN selection regime

The experiments reported in this study were carried out using the EPN populations, consisting of twelve populations categorized into three selection regimes (Singh et al 2021).

#### E_1-4_: Populations selected for better survival following infection with the Gram-positive bacterium *Enterococcus faecalis*

Every generation, 2–3-day old adult flies (200 females and 200 males) are subjected to infection with *E. faecalis*, and 96-hours post-infection, the survivors are allowed to reproduce and contribute to the next generation. At the end of 96 hours, on average 100 females and 100 males are left alive in each of the E_1-4_ populations.

#### P_1-4_: Sham-infected control populations

Every generation, 2–3-day old adult flies (100 females and 100 males) are subjected to sham-infection, and 96-hours post-sham-infection, the survivors are allowed to reproduce and contribute to the next generation.

#### N_1-4_: Uninfected control populations

Every generation, 2–3-day old adult flies (100 females and 100 males) are subjected to light CO_2_ anesthesia only, and 96-hours post-procedure, the survivors are allowed to reproduce and contribute to the next generation. (Under usual circumstances, no mortality occurs in the P_1-4_ and N_1-4_ populations during maintenance of the selection regimes.)

The EPN populations were derived from the ancestral Blue Ridge Baseline (BRB_1-4_) populations. The E_1_, P_1_, and N_1_ populations were derived from the BRB_1_ population and constitutes ‘block 1’ of the experimental evolution regime. Similarly, E_2_, P_2_, and N_2_ populations were derived from the BRB_2_ population and constitutes ‘block 2’, and so on. This *block design* implies that E_1_, P_1_, and N_1_ have a more recent common ancestor, compared to E_1_ and E_2_, or P_1_ and P_2_, and so on. Populations belonging to each block were handled together, both during maintenance of the populations and during experiments. Blocks were also used as experimental and statistical replicates.

The maintenance of the EPN populations have been previously described (Singh et al 2021, Singh et al 2022). The EPN populations are maintained on banana-jaggery-yeast food medium. Every generation, eggs are collected at a density of 60-80 eggs per vial (with 6-8 ml food medium). 10 such rearing vials (9 cm height × 2.5 cm diameter) are set up for each of the 12 populations. These vials are incubated at 25 °C, 60% RH, and a 12:12 LD cycle. Under such maintenance conditions, eggs develop into adults within 9-10 days of egg collection. On day 12 post-egg laying (PEL), flies from each population are subjected to selection according to their identity, as described above. The adults stay in the rearing vial till day 12 PEL, and are sexually mature, and sexually active, by the time they are subjected to selection. After being subjected to selection, the flies are housed in plexiglass cages (14 cm × 16 cm × 13 cm), one cage for each population. The cages are provided with fresh food medium, on a 60 mm Petri plate, on every alternate day. On day 16 PEL eggs are collected from the surviving flies in each cage to start the next generation.

### 2.3 Pre-experiment standardization

Prior to experiments, flies from the three selection regimes were reared for a generation under ancestral maintenance conditions. This is done to account for any non-genetic parental effects (Rose 1984), and flies thus generated are referred to as *standardized* flies. To generate standardized flies, eggs were collected from all the populations at a density of 60-80 eggs per vial; 10 such vials were set up per population. The vials were incubated under standard maintenance conditions described above. On day 12 PEL, the adults were transferred to plexiglass cages (14 cm × 16 cm × 13 cm) with food plates (Petri plates, 60 mm diameter). Eggs for experimental flies were collected from these *standardised* population cages.

### 2.4 Experiment design

#### Experiment 1.a. Effect of mating on post-infection survival of females from E, P, and N populations when infected with the native pathogen (*E. faecalis*)

In this experiment we tested if focal females from E, P, and N populations exhibited mating-induced increase in susceptibility to *E. faecalis* when mated with common BRB males. The experiment for each block was carried out separately. This experiment was done after 45 generations of forward selection.

Eggs were collected from standardized E, P, and N flies, at a density of 60-80 eggs per vial; 20 such vials were set up per population. Similarly, eggs were collected from the BRB flies, at a density of 60-80 eggs per vial; 30 such vials were set up. These vials were incubated under standard maintenance conditions (described in section 2.2), and on 10^th^ day post-egg laying (PEL), freshly eclosing flies were collected as virgins and housed in single sex vials. Virgin females were collected for the E, P, and N populations, and housed at a density of 8 females per vial (each vial with 1.5-2 ml of standard food medium); 50 vials of virgin females were collected per population. Virgin males were collected from BRB population and housed at a density of 10 males per vial; 90 vials of virgin males were collected.

On 12^th^ day PEL, to obtain mated females, 30 vials of virgin females from each population (E, P, and N) were combined (without anesthesia) individually with vials of BRB virgin males, individually in fresh food vials. These vials were visually observed to ensure that the each of the eight females in a vial had mated at least once. Thereafter, 20 vials from each population were set aside for infection, and the other 10 vials were monitored for re-mating rate (see below). The females and males continued to be housed together from the point of initiation of the mating set-up till the time of infection. The remaining 20 vials of virgin females from each population (E, P, and N) were simply transferred to fresh food vials. 4-5 hours after the initiation of the mating set-up, mated females and males were anesthetized, vial by vial, and the females were subjected to infection with *E. faecalis* (or sham-infections); the males were discarded. Virgin females were also infected simultaneously. After infections, the females were placed in fresh food vials. In total, 10 vials of infected females (n = 80 females) and 5 vials of sham-infected females (n = 40 females) were set up per population (E, P, and N), per mating status (virgin and mated), per block. These vials were monitored for mortality, every 4-6 hours, for 96 hours post-infection. Flies alive at the end of 48 hours were shifted to fresh food vials, and flies alive at the end of 96 hours were discarded (right censored).

To get an estimate of re-mating rate of females form different populations, 10 vials per population (E, P, and N) were monitored, every 15 minutes, for 5 hours. The total number of mating events were recorded for each vial. The number of mating events from each vial was used as the unit of replication for remating rate. (Since all vials had the same number of females and males, the absolute number of mating events was used for analysis without any per-female normalization.)

#### Experiment 1.b. Effect of mating on post-infection survival of males from E, P, and N populations when infected with the native pathogen (*E. faecalis*)

In this experiment we tested if focal males from E, P, and N populations exhibited mating-induced increase in susceptibility to *E. faecalis* when mated with common BRB females. The experiment for each block was carried out separately. This experiment was done after 45 generations of forward selection.

Eggs were collected from standardized E, P, and N flies, at a density of 60-80 eggs per vial; 20 such vials were set up per population. Similarly, eggs were collected from the BRB flies, at a density of 60-80 eggs per vial; 30 such vials were set up. These vials were incubated under standard maintenance conditions (described in section 2.2), and on 10^th^ day post-egg laying (PEL), freshly eclosing flies were collected as virgins and housed in single sex vials. Virgin males were collected for the E, P, and N populations, and housed at a density of 8 males per vial (each vial with 1.5-2 ml of standard food medium); 50 vials of virgin males were collected per population. Virgin females were collected from BRB population and housed at a density of 10 females per vial; 90 vials of virgin females were collected.

On 12^th^ day PEL, to obtain mated males, 30 vials of virgin males from each population (E, P, and N) were combined (without anesthesia) individually with vials of BRB virgin females, individually in fresh food vials. These vials were visually observed to ensure that the each of the eight males in a vial had mated at least once. Thereafter, 20 vials from each population were set aside for infection, and the other 10 vials were monitored for re-mating rate (see below). The males and females continued to be housed together from the point of initiation of the mating set-up till the time of infection. The remaining 20 vials of virgin males from each population (E, P, and N) were simply transferred to fresh food vials. 4-5 hours after the initiation of the mating set-up, mated males and females were anesthetized, vial by vial, and the males were subjected to infection with *E. faecalis* (or sham-infections); the females were discarded. Virgin males were also infected simultaneously. After infections, the males were placed in fresh food vials. In total, 10 vials of infected males (n = 80 males) and 5 vials of sham-infected males (n = 40 males) were set up per population (E, P, and N), per mating status (virgin and mated), per block. These vials were monitored for mortality, every 4-6 hours, for 96 hours post-infection. Flies alive at the end of 48 hours were shifted to fresh food vials, and flies alive at the end of 96 hours were discarded (right censored).

To get an estimate of re-mating rate of males form different populations, 10 vials per population (E, P, and N) were monitored, every 15 minutes, for 5 hours. The total number of mating events were recorded for each vial. The number of mating events from each vial was used as the unit of replication for remating rate. (Since all vials had the same number of males and females, the absolute number of mating events was used for analysis without any per-male normalization.)

#### Experiment 2. Effect of mating on post-infection survival of females from E and P populations when infected with two novel pathogens (*B. thuringiensis* and *P. entomophila*)

In this experiment we tested if focal females from E and P populations exhibited mating-induced increase in susceptibility to *B. thuringiensis* and *P. entomophila* when mated with common BRB males. The experiment for each block was carried out separately. This experiment was done after 55 generations of forward selection.

Eggs were collected from standardized E and P flies, at a density of 60-80 eggs per vial; 20 such vials were set up per population. Similarly, eggs were collected from the BRB flies, at a density of 60-80 eggs per vial; 25 such vials were set up. These vials were incubated under standard maintenance conditions (described in section 2.2), and on 10^th^ day post-egg laying (PEL), freshly eclosing flies were collected as virgins and housed in single sex vials. Virgin females were collected for the E and P populations, and housed at a density of 8 females per vial (each vial with 1.5-2 ml of standard food medium); 50 vials of virgin females were collected per population. Virgin males were collected from BRB population and housed at a density of 10 males per vial; 75 vials of virgin males were collected.

On 12^th^ day PEL, to obtain mated females, 40 vials of virgin females from each population (E and P) were combined (without anesthesia) individually with vials of BRB virgin males, individually in fresh food vials. These vials were visually observed to ensure that the each of the eight females in a vial had mated at least once. The females and males continued to be housed together from the point of initiation of the mating set-up till the time of infection. The remaining 20 vials of virgin females from each population (E and P) were simply transferred to fresh food vials. 4-5 hours after the initiation of the mating set-up, mated females and males were anesthetized, vial by vial, and the females were subjected to infection with either *B. thuringiensis* or *P. entomophila* (or sham-infections); the males were discarded. Virgin females were also infected simultaneously. After infections, the females were placed in fresh food vials. In total, 10 vials of *B. thuringiensis* infected females (n = 80 females), 10 vials of *P. entomophila* infected females (n = 80 females), and 5 vials of sham-infected females (n = 40 females) were set up per population (E and P), per mating status (virgin and mated), per block. These vials were monitored for mortality, every 4-6 hours, for 96 hours post-infection. Flies alive at the end of 48 hours were shifted to fresh food vials, and flies alive at the end of 96 hours were discarded (right censored).

### 2.5 Statistical analysis

Survival data of infected flies, from experiments 1(a), 1(b), and 2, was modeled as Survival ∽ Selection history + Mating status + (Selection history × Mating status) + (1|Block), using mixed-effects Cox proportional hazards, where selection history, mating status, and their interaction were modeled as fixed factors, and block identity was modeled as a random factor. This model was subjected to analysis of deviance (type II) for significance testing for the fixed factors (tabulated in Table 1). Survival data for the three pathogens, and both sexes, were analyzed separately. Since there was negligible mortality observed in sham-infected flies (Figure 1), only the survival data of infected flies were subjected to statistical analysis.

**Table 1.** Analysis of deviance (type II) for the effect of selection history, mating status, and their interaction on post-infection survival of virgin and mated flies from the E, P, and N populations: (a) females infected with *Enterococcus faecalis*; (b) males infected with *Enterococcus faecalis*; (c) females infected with *Bacillus thuringiensis*; and (d) females infected with *Pseudomonas entomophila*.

| Fixed factors | DF | Chi-square value | p-value |
| --- | --- | --- | --- |
| (a) Survival of E, P, and N females, either virgin or mated, when infected with *Enterococcus faecalis* |  |  |  |
| **Selection history** | **2** | **133.8540** | **<2.2e-16** |
| Mating status | 1 | 0.4964 | 0.4811 |
| Selection history × Mating status | 2 | 2.3661 | 0.3063 |
| (b) Survival of E, P, and N males, either virgin or mated, when infected with *Enterococcus faecalis* |  |  |  |
| **Selection history** | **2** | **56.2585** | **6.076e-13** |
| Mating status | 1 | 0.3799 | 0.5377 |
| Selection history × Mating status | 2 | 0.1319 | 0.9362 |
| (c) Survival of E and P females, either virgin or mated, when infected with *Bacillus thuringiensis* |  |  |  |
| **Selection history** | **1** | **224.067** | **<2.2e-16** |
| **Mating status** | **1** | **16.080** | **6.073e-05** |
| **Selection history × Mating status** | **1** | **17.146** | **3.461e-05** |
| (d) Survival of E and P females, either virgin or mated, when infected with *Pseudomonas entomophila* |  |  |  |
| **Selection history** | **1** | **267.3126** | **<2.2e-16** |
| **Mating status** | **1** | **27.7184** | **1.403e-07** |
| **Selection history × Mating status** | **1** | **5.6996** | **0.01697** |

**Figure 1.**
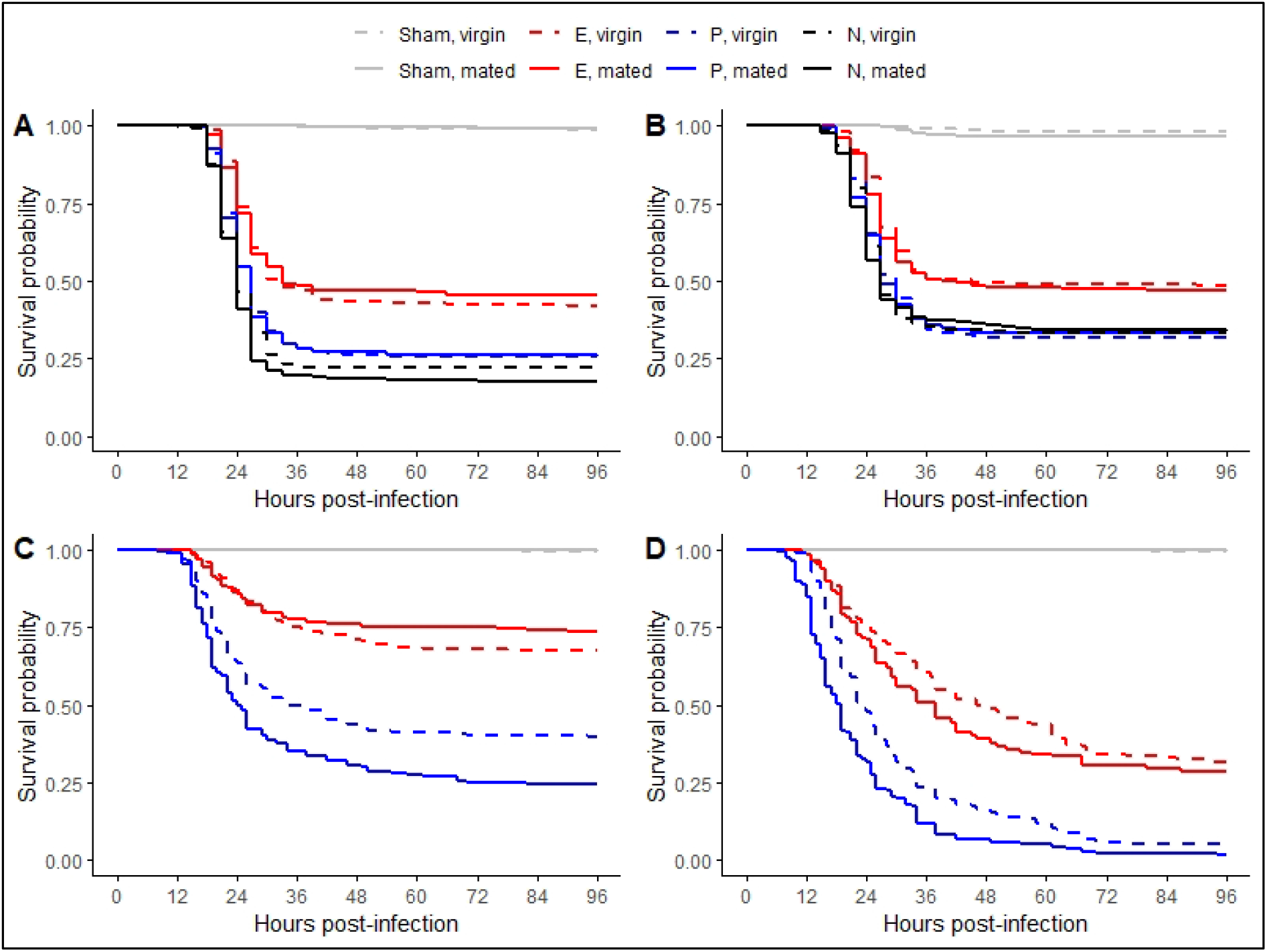
Post-infection survival of virgin and mated flies from the E, P, and N populations: (A) females infected with *Enterococcus faecalis*; (B) males infected with *Enterococcus faecalis*; (C) females infected with *Bacillus thuringiensis*; and (D) females infected with *Pseudomonas entomophila*.

Hazard ratio for mortality of infected mated flies, relative to the infected virgin flies, was calculated for flies from each population (E, P, and N) using the mixed-effects Cox-proportional hazards model

Survival ∽ Mating status + (1|Block),

where mating status was used as a fixed factor and block identity as a random factor. Hazard ratios for the three pathogens, and both sexes, were calculated separately (represented graphically in Figure 2, and tabulated in Table S1).

**Figure 2.**
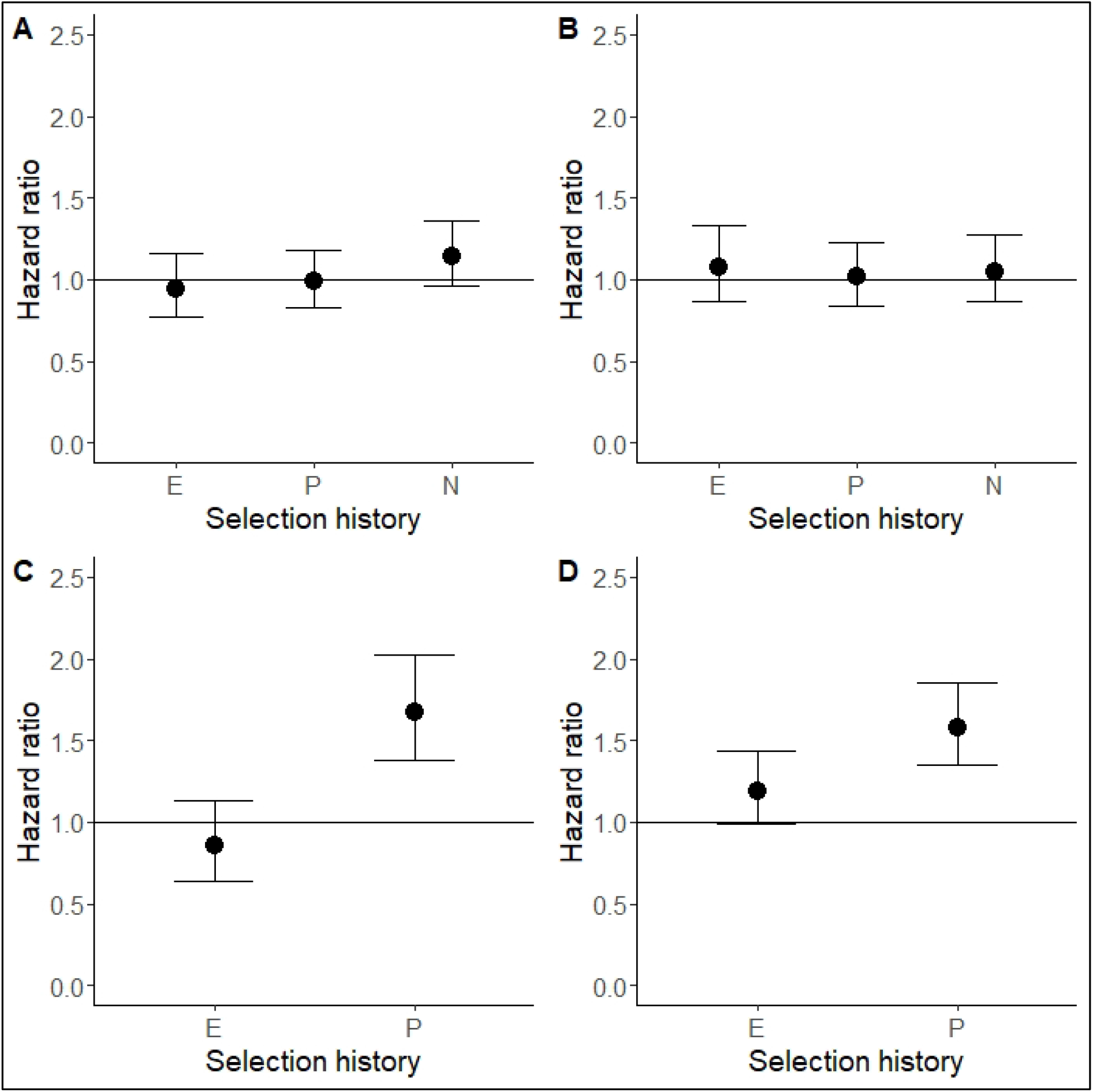
Hazard ratio (mixed-effects Cox proportional hazards method) for mortality of mated flies, relative to virgin flies, from the E, P, and N populations: (A) females infected with *Enterococcus faecalis*; (B) males infected with *Enterococcus faecalis*; (C) females infected with *Bacillus thuringiensis*; and (D) females infected with *Pseudomonas entomophila*. The horizontal line, at hazard ratio equals to 1, represents the hazard for virgin female for the corresponding population. (Error bars represent 95% confidence intervals.)

Remating rate data was analyzed using analysis of variance (ANOVA, type-III) where selection history was included as a fixed factor, and block identity and selection × block interaction were included as random effects. Significance tests for the random effects included in ANOVA are tabulated in Table S2.

All analyses were carried out using R statistical software (version 4.1.0; R Core Team 2021), using various functions from the *survival* (Therneau 2021), *coxme* (Therneau 2020), and *lmerTest* (Kuznetsova et al 2017) packages. Graphs were created using the *ggplot2* (Wikham 2016) and *survminer* (Kassambara et al 2021) packages.

## 3. Results

### 3.1 Survival of females infected with *Enterococcus faecalis*

Post-infection survival of females when infected with *E. faecalis* was significantly affected by selection history, but not by mating status, or selection history × mating status interaction (Table 1.a, Figure 1.a). The mated females from either E (hazard ratio, 95% confidence interval: 0.945, 0.769-1.161), P (HR, 95% CI: 0.989, 0.826-1.183), or N (HR, 95% CI: 1.143, 0.962-1.360) population did not differ in post-infection survival relative to the virgin females (Figure 2.a).

### 3.2 Survival of males infected with *E. faecalis*

Post-infection survival of males when infected with *E. faecalis* was significantly affected by selection history, but not by mating status, or selection history × mating status interaction (Table 1.b, Figure 1.b). The mated males from either E (HR, 95% CI: 1.074, 0.866-1.332), P (HR, 95% CI: 1.013, 0.839-1.224), or N (HR, 95% CI: 1.049, 0.866-1.270) population did not differ in post-infection survival relative to the virgin males (Figure 2.b).

### 3.3 Survival of females infected with *Bacillus thuringiensis*

Post-infection survival of females when infected with *B. thuringiensis* was significantly affected by selection history, mating status, and selection history × mating status interaction (Table 1.c, Figure 1.c). Pooling females from both mating treatments, E females (HR, 95% CI: 0.266, 0.224-0.317) survived better relative to P females. Among E females, there was no difference between survival of virgin and mated females (HR, 95% CI: 0.852, 0.640-1.135), but among P females, mated females (HR, 95% CI: 1.673, 1.383-2.025) exhibited greater mortality relative to virgin females (Figure 2.c).

### 3.4 Survival of females infected with *Pseudomonas entomophila*

Post-infection survival of females when infected with *P. entomophila* was significantly affected by selection history, mating status, and selection history × mating status interaction (Table 1.d, Figure 1.d). Pooling females from both mating treatments, E females (HR, 95% CI: 0.355, 0.313-0.403) survived better relative to P females. Among E females, there was no difference between survival of virgin and mated females (HR, 95% CI: 1.191, 0.989-1.434), but among P females, mated females (HR, 95% CI: 1.582, 1.349-1.854) exhibited greater mortality relative to virgin females (Figure 2.d).

### 3.5 Remating rate

Remating rate was not affected by selection history in case of either female (F_2,8_ = 1.222, p = 0.344) or male (F_2,8_ = 0.36, p = 0.708) flies from the E, P, and N populations (Figure 3).

**Figure 3.**
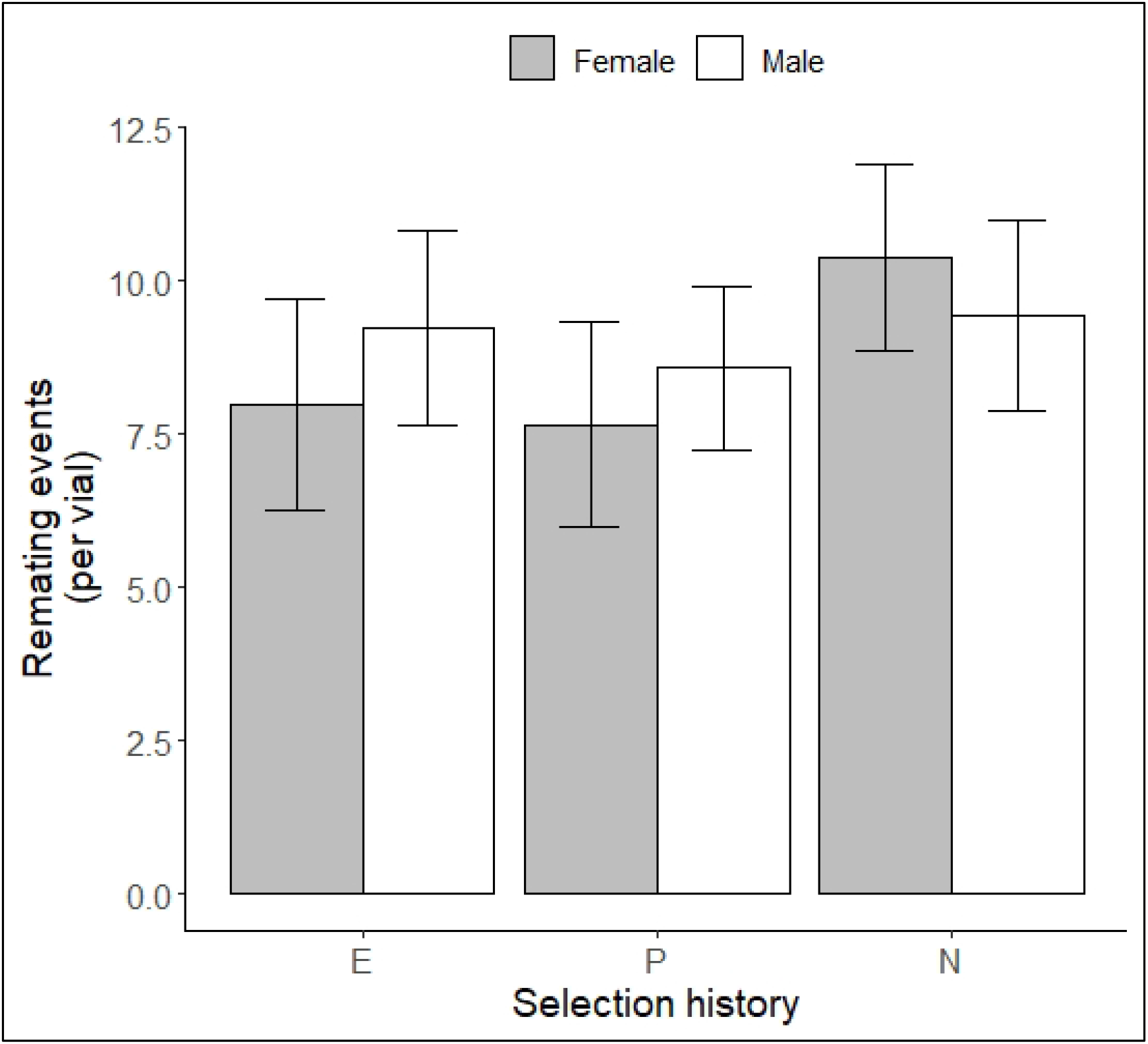
Remating rate of females and males of the E, P, and N populations. (Error bars represent 95% confidence intervals.)

## 4. Discussion and Conclusion

In this study, we measured the effect of sexual activity (mating) on post-infection survival of female and male flies (*Drosophila melanogaster*) upon infection with bacterial pathogens. We tested if the effect of mating on post-infection survival was determined by the evolutionary history of the hosts. We experimentally evolved fly populations for increased survival following infection with a Gram-positive bacterium, *Enterococcus faecalis* (Singh et al 2021). We infected virgin and mated females from these evolved (E_1-4_) populations, and their ancestrally paired control (P_1-4_ and N_1-4_) populations (see Materials and Methods for more details), with three bacterial pathogens: *E. faecalis, Bacillus thuringiensis*, and *Pseudomonas entomophila. E. faecalis* is the *native* pathogen, i.e., the pathogen used for selection in the experimental evolution set-up. *B. thuringiensis* (Gram-positive) and *P. entomophila* (Gram-negative) represent two *novel* pathogens, one of which has the same Gram-character as the native pathogen while the other doesn’t.

When infected with *E. faecalis*, the females from the selected populations had better survival compared to the females from the control populations (figure 1.a). This proved that the selection process was a success, and this observation agreed with results reported previously (Singh et al 2021, Singh et al 2022). Additionally, the mated and the virgin females did not differ from one another in terms of post infection survival when infected with *E. faecalis* (figure 2.a). When infected with the novel pathogens, the females from the selected populations again survived better than the females of the control populations, in case of both *B. thuringiensis* (figure 1.c) and *P. entomophila* (figure 1.d). This replicated and verified previous results from these populations where it was demonstrated that the selected populations have evolved cross-resistance against a wide range of novel bacterial pathogens (Singh et al 2021). Furthermore, in case of both *B. thuringiensis* (figure 2.c) and *P. entomophila* (figure 2.d), the mated females from the control populations died more after being infected, relative to the virgin females. There was no difference in post-infection survival of virgin and mated females from the selected populations in case of either of the novel pathogens (figure 2.c and 2.d).

Altogether, results from our experiments suggest that females from the selected populations do not suffer any mating induced change in susceptibility to bacterial infections. On the other hand, females from the control populations do suffer a mating induced increase in susceptibility to bacterial infections, but in a pathogen specific manner. Pathogen specific differences in survival between virgin and mated females has been reported previously in other studies (Short and Lazzaro 2010, Basu et al 2022). We report here that the pathogen specificity of the mating-induced change in susceptibility to infections can be modified by host evolutionary history. Since we did not find any difference in re-mating rate of females from the selected and the control populations (figure 3), the differences in mating-induced increase in susceptibility to infections cannot be explained by differential re-mating rate. Previous studies have also shown that mating rate has no effect on mating-induced immune suppression in females but does have a major effect in case of males (McKean and Nunney 2001, McKean and Nunney 2005).

When we infected virgin and mated males from the selected and the control populations with *E. faecalis*, males from selected populations in general survived better than the males from control populations (figure 1.b), attesting to that fact that selection had successfully worked in the E populations (Singh et al 2021), but post-infection survival was not affected by mating in any of the populations (figure 2.b). Since we only infected males with the native pathogen, we cannot comment on either evolution of cross-resistance or pathogen-specific mating induced change in susceptibility to pathogens. In case of males too, the re-mating rate did not differ between the selected and the control populations (figure 3).

During regular maintenance of the experimental evolution regime, the flies of the selected (E_1-4_) populations are already mated at the time of infection. Therefore, we hypothesized that E flies must be under selection to evolve to counteract the negative effects of mating on immune function, if there are indeed any negative effects. Our results show that E females are in fact better at counteracting the negative effects of mating on immune function (measured as post-infection survival). This may be driven by either (a) E females having evolved to optimize immune defense against bacterial pathogens with a resource deprived immune system (because of mating-induced re-routing of resources towards reproduction), or (b) E females having evolved to counteract the mating-dependent signals that encourage females to invest towards reproduction. Based on the present data we cannot differentiate between these two possibilities. In *D. melanogaster* females, sex peptide transferred from males during mating induces production of Juvenile Hormone (JH) in female corpus allota, which reduces a female’s ability to defend against bacterial infections (Schwenke and Lazzaro 2017). JH also encourages investment towards reproductive functions (Flatt et al 2005) and suppresses expression of AMP genes (Flatt et al 2008). Therefore, we hypothesize that the selected populations may have evolved to be resistant to JH-mediated modifications of the immune system that follow sexual activity in females.

It has been previously suggested that starvation induces a re-structuring of the insect immune system towards a new functional equilibrium, so that the immune system can maintain optimal functionality even in a resource limited environment (Adamo et al 2016, Adamo 2017). Based on the results obtained in this study, and in previously published studies (discussed in the Introduction), we hypothesize that sexual activity (mating) has a similar influence on the insect immune system, and that this influence is specific to each host species. This would explain (a) why different immune components are affected differently by mating (viz. Fedorka et al 2004, Castella et al 2009), (b) why the same immune components are affected differently in different species (viz. Fedorka et al 2004, Rolf and Siva-Jothy 2002, Castella et al 2009), and (c) why the effect of mating on post-infection survival is pathogen specific (Short and Lazzaro 2010, Basu et al 2022). The exact mechanism governing this mating-induced re-structuring of insect immune system is yet to be defined, but it can be assumed that Juvenile Hormone (JH) is a key regulator of this process. In *D. melanogaster*, increased production of JH in mated females, induced by sex peptide transferred by males during copulation, increases susceptibility of females to bacterial infections (Schwenke and Lazzaro 2017). Inhibition of JH synthesis nullifies the negative effects of mating on PO activity in male and female *T. molitor* (Rolf and Siva-Jothy 2002). Additionally, signals from the germline cells may also be key regulators of mating-induced re-structuring of the insect immune system (Fedorka et al 2007, Short et al 2012, Short and Lazzaro 2013, Rodrigues et al 2021).

Assuming our above hypothesis holds true, we further propose that the evolutionary history of the host organism decides the degree of re-structuring that occurs in a mated individual, compared to a virgin individual. This would explain why females from our selected populations do not exhibit any mating-induced change in susceptibility to pathogens, both native and novel, but the females from the control populations do exhibit increased mortality after mating in case of both novel pathogens. It is possible that host sex also determines the degree of immune system re-structuring, but we are not able to comment on that because we only measured post-infection mortality of the males using the native pathogen.

To summarize, using fly populations experimentally evolved to better survive bacterial (*E. faecalis*) infections, we tested the role of host evolutionary history in determining if sexually active flies are more susceptible to bacterial infection compared to sexually inactive flies. We observe that whether mated females are more susceptible to infection is dependent upon the pathogen used for infection. In case of pathogens where we do observe an increased susceptibility of mated females to infection, females from the evolved populations exhibit no mating-induced change in susceptibility to infections, while the females from the control populations become more susceptible to infections after mating. This suggests that host evolutionary history, and adaptation to better survive a pathogen challenge, reduces a host’s vulnerability to mating-induced immune suppression in *Drosophila melanogaster*.

## Acknowledgements

The authors thank Prof. P Cornelis (Vrije Universiteit Brussel, Belgium) for providing us with *Pseudomonas entomophila* L48, and Prof. B Lazzaro (Cornell University, USA) for providing us with *Enterococcus faecalis* isolate.

## Supplementary Materials

**Table S1**. Hazard ratio of mated flies relative to virgin flies from E, P, and N populations, when subjected to infection with different pathogens.

**Table S2**. Significance tests for random effect included in analysis of variance (ANOVA) of remating rate data for (a) females, and (b) males.

## Notes

Funding: The study was funded by intramural funding from IISER Mohali, India. AB was supported by Senior Research Fellowship for graduate students from CSIR, Govt. of India. AS was supported by Senior Research Fellowship for graduate students from University Grants Commission, Govt. of India. RBG was supported by Summer Research Fellowship Program for undergraduate students funded by JNCASR, Bangalore.

### Competing Interest Statement

The authors have declared no competing interest.

