## Supplemental Table 1 for "Selection for increased post-infection survival ameliorates mating induced immune suppression in *Drosophila melanogaster* females"

Table S1. Hazard ratio of mated flies relative to virgin flies from E, P, and N populations, when subjected to infection with different pathogens.

| Pathogen | Sex | Population | Reference  treatment | Focal  treatment | HR | Lower  CI | Upper  CI | Z | p-value | Random  factor | Random  factor  variance |
| --- | --- | --- | --- | --- | --- | --- | --- | --- | --- | --- | --- |
| *E. faecalis* | Female | E | Virgin | Mated | 0.9448002 | 0.7685727 | 1.161435 | -0.54 | 0.59 | Block | 0.000112372 |
| *E. faecalis* | Female | P | Virgin | Mated | 0.9886988 | 0.8259657 | 1.183494 | -0.12 | 0.9 | Block | 0.04119653 |
| *E. faecalis* | Female | N | Virgin | Mated | 1.14349 | 0.9616199 | 1.359758 | 1.52 | 0.13 | Block | 0.0183106 |
| *E. faecalis* | Male | E | Virgin | Mated | 1.07417 | 0.8664926 | 1.331622 | 0.65 | 0.51 | Block | 0.000829956 |
| *E. faecalis* | Male | P | Virgin | Mated | 1.013532 | 0.8392306 | 1.224035 | 0.14 | 0.89 | Block | 0.009754843 |
| *E. faecalis* | Male | N | Virgin | Mated | 1.048747 | 0.8662518 | 1.269689 | 0.49 | 0.63 | Block | 0.06643679 |
| *B. thuringiensis* | Female | E | Virgin | Mated | 0.8520461 | 0.6395775 | 1.135097 | -1.09 | 0.27 | Block | 0.9320261 |
| *B. thuringiensis* | Female | P | Virgin | Mated | 1.67337 | 1.382942 | 2.02479 | 5.29 | 1.20E-07 | Block | 0.4380157 |
| *P. entomophila* | Female | E | Virgin | Mated | 1.191116 | 0.989132 | 1.434345 | 1.84 | 0.065 | Block | 0.2798338 |
| *P. entomophila* | Female | P | Virgin | Mated | 1.581516 | 1.348712 | 1.854504 | 5.64 | 1.70E-08 | Block | 0.06133306 |
