## Supplemental Table 2 for "Selection for increased post-infection survival ameliorates mating induced immune suppression in *Drosophila melanogaster* females"

Table S2. Significance tests for random effect included in analysis of variance (ANOVA) of remating data for (a) females, and (b) males.

|  | Number of parameters | Log-likelihood | AIC | LRT | DF | p-value (Chi-square) |
| --- | --- | --- | --- | --- | --- | --- |
| (a) Female remating rate | | | | | | |
| <none> | 6 | -358.79 | 729.58 |  |  |  |
| (1\|Block) | 5 | -358.79 | 727.59 | 0.0062 | 1 | 0.93700 |
| (1\|Block:Population) | 5 | -363.16 | 736.31 | 8.7301 | 1 | 0.00313 |
| (b) Male remating rate | | | | | | |
| <none> | 6 | -332.23 | 676.47 |  |  |  |
| (1\|Block) | 5 | -336.24 | 682.47 | 8.0061 | 1 | 0.004662 |
| (1\|Block:Population) | 5 | -332.82 | 675.64 | 1.1771 | 1 | 0.277950 |
